# Inhibiting SARS-CoV-2 infection *in vitro* by suppressing its receptor, angiotensin-converting enzyme 2, via aryl-hydrocarbon receptor signal

**DOI:** 10.1101/2021.03.04.433658

**Authors:** Keiji Tanimoto, Kiichi Hirota, Takahiro Fukazawa, Yoshiyuki Matsuo, Toshihito Nomura, Nazmul Tanuza, Nobuyuki Hirohashi, Hidemasa Bono, Takemasa Sakaguchi

## Abstract

Since understanding molecular mechanisms of SARS-CoV-2 infection is extremely important for developing effective therapies against COVID-19, we focused on the internalization mechanism of SARS-CoV-2 via ACE2. Although cigarette smoke is generally believed to be harmful to the pathogenesis of COVID-19, cigarette smoke extract (CSE) treatments were surprisingly found to suppress the expression of ACE2 in HepG2 cells. We thus tried to clarify the mechanism of CSE effects on expression of ACE2 in mammalian cells. Because RNA-seq analysis suggested that suppressive effects on *ACE2* might be inversely correlated with induction of the genes regulated by aryl hydrocarbon receptor (AHR), the AHR agonists 6-formylindolo(3,2-b)carbazole (FICZ) and omeprazole (OMP) were tested to assess whether those treatments affected ACE2 expression. Both FICZ and OMP clearly suppressed *ACE2* expression in a dose-dependent manner along with inducing *CYP1A1*. Knock-down experiments indicated a reduction of *ACE2* by FICZ treatment in an AHR-dependent manner. Finally, treatments of AHR agonists inhibited SARS-CoV-2 infection into Vero E6 cells as determined with immunoblotting analyses detecting SARS-CoV-2 specific nucleocapsid protein. We here demonstrate that treatment with AHR agonists, including CSE, FICZ, and OMP, decreases expression of ACE2 via AHR activation, resulting in suppression of SARS-CoV-2 infection in mammalian cells.

## Introduction

Severe acute respiratory syndrome coronavirus 2 (SARS-CoV-2), which was first reported in Wuhan, China in December 2019, is rapidly and continuously spreading all over the world [World Health Organization (WHO)/Coronavirus disease (SARS-CoV-2 induced disease: COVID-19) pandemic] (WHO website). Many epidemiological studies have suggested that smoking habits have serious effects on the pathogenesis of COVID-19 (Vardavas et al., 2020; Berlin et al., 2020; Guo, 2020; Kashyap, 2020). Expression of the receptor for SARS-CoV-2 infection, angiotensin-converting enzyme 2 (ACE2), was also reported to be higher in smoking mice and humans, suggesting that smokers may be at a higher risk of infection (Smith et al., 2020). On the other hand, several reports suggested fewer smokers among patients infected with SARS-CoV-2 or lower numbers of SARS-CoV-2 positive cases among smokers than among non-smokers (Rentsch et al, 2020; Miyara et al, 2020; Lusignan et al,2020; Williamson et al., 2020). Therefore, the impact of smoking on SARS-CoV-2 infection is unclear. Cigarette smoke contains a variety of compounds, such as polycyclic aromatic hydrocarbons (PAHs) and nitrosamines, and cellular responses to cigarette smoke exposure are thus diverse (Zhang et al., 2012; AlQasrawi et al., 2020). PAHs bind to and activate aryl hydrocarbon receptor (AHR, which is an intracellular receptor type transcription factor), and induce various cellular physiological and pathological responses through the regulation of gene expression (Hankinson, 1995; Moorthy et al., 2015). AHR is also activated by a broad variety of exogenous and endogenous small molecular weight compounds, such as 2,3,7,8-tetrachlorodibenzo-*p*-dioxin (TCDD), 3,3’,4,4’,5-pentachlorobiphenyl-126 (PCB126), 6-formylindolo(3,2-b)carbazole (FICZ), and omeprazole, and their physiological and pathological significance, especially in regard to the immune system, have received much attention (Denison et al., 2003; Gutiérrez-Vázquez et al., 2018; Rothhammer et al., 2019).

In this study, we therefore aimed to clarify the effects of components of cigarette smoke and AHR agonists on ACE2 expression in mammalian cells to better understand the role of ACE2 in the SARS-CoV-2 internalization mechanism.

## Results

### Effects of cigarette smoke extract (CSE) on *ACE2* expression in human cell lines

At first we examined expression levels of the SARS-CoV-2 viral receptor ACE2 in various cell lines by using quantitative RT-PCR (qRT-PCR) (Supplementary File 1a). Results demonstrated that *ACE2* expression level varied among the cell lines, and HSC2 (oral cavity origin), PC9 (lung origin), and HepG2 (liver origin), which had the highest levels of *ACE2* expression, were used in the first experiment. The cells were treated with various doses of cigarette smoke extract (CSE) for 24 hours, after which expression level of cytochrome P450 family 1 subfamily A member 1 (*CYP1A1*) gene, which is known to be CSE-inducible, was evaluated with qRT-PCR. As expected, CSE treatment induced expression of *CYP1A1* in HepG2 cells in a dose-dependent manner (Figure 1a). CSE treatment also induced expression of *CYP1A1* in PC9 cells, but expression was only slightly increased in HSC2 cells (Supplementary File 1b). Interestingly, *ACE2* expression was significantly reduced in CSE-treated HepG2 cells in a dose-dependent manner, and was slightly reduced in PC9 and HSC2 cells (Figure 1b and Supplementary File 1c).

**Figure 1.**
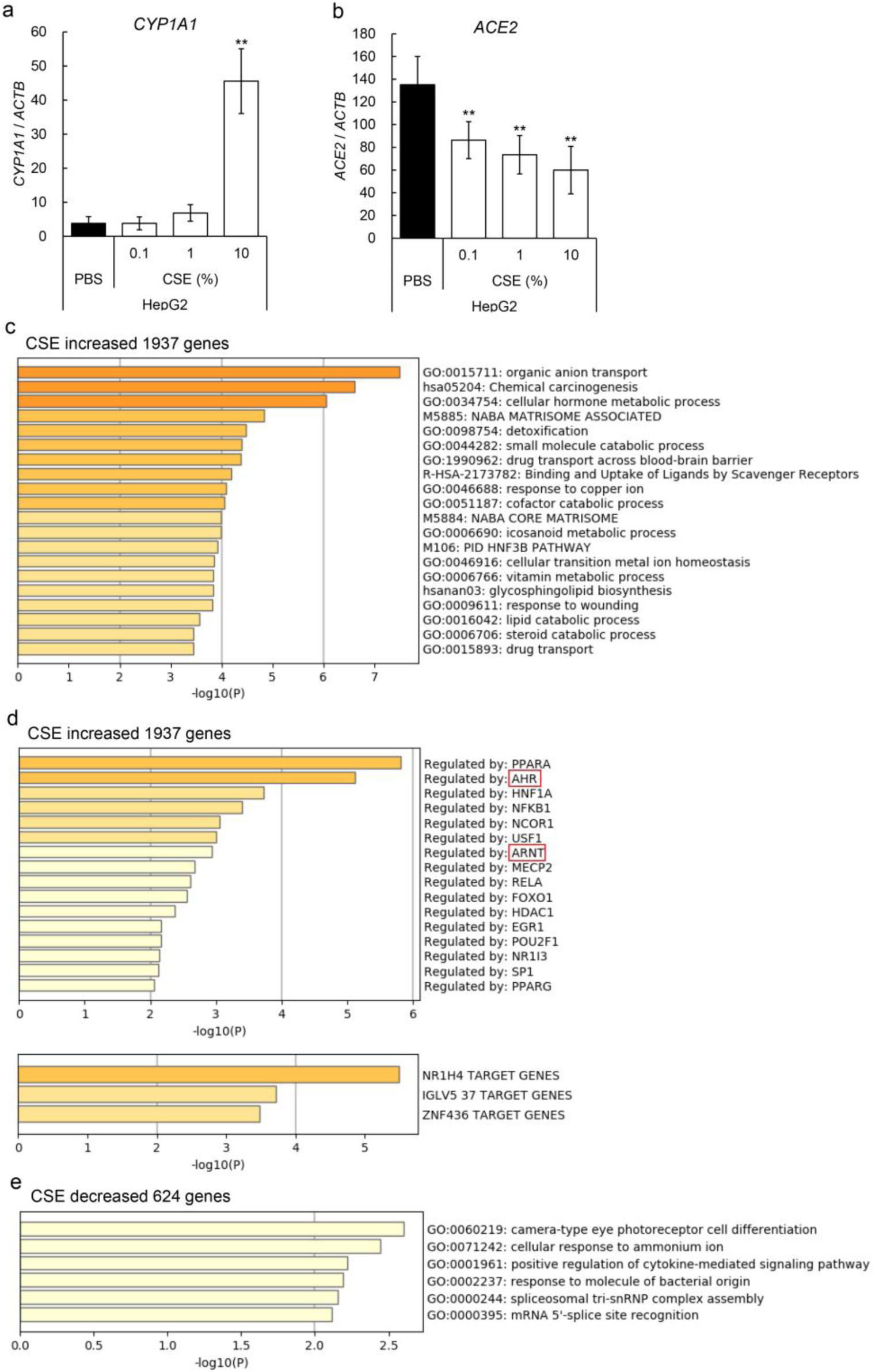
Effects of CSE on *ACE2* expression in human cell lines. (a, b) Expression levels of *CYP1A1* and *ACE2* genes in various concentrations of CSE treated HepG2 (Liver origin) cells for 24 hours were evaluated by qRT-PCR. Relative gene expression levels were calculated by using *ACTB* expression as the denominator for each cell line (n = 3). For all quantitative values, the average and SD are shown. Statistical significance was calculated for the indicated paired samples with ** representing *P* < 0.01. Gene set enrichment analyses for gene lists analyzed by RNA-seq were performed with Metascape. Upregulated genes [log_2_(CSE-DMSO) ≥ 1] (c) in CSE treated HepG2 cells and downregulated genes [log_2_(CSE-DMSO) ≤ −1] (e) were enriched by gene ontology (GO) term. Process enrichment analyses using TRRUST showed human transcriptional regulatory interactions in upregulated genes (d) and downregulated genes (f). AHR and ARNT, hetero-dimeric transcriptional partners, are indicated with red boxes.

### RNA-seq analysis of CSE treated HepG2 cells

To elucidate cellular responses to treatment with CSE, comprehensive gene expression was investigated with RNA-seq analysis. Results demonstrated that CSE increased the expression of 1937 genes and decreased that of 624 genes in HepG2 cells. Gene set enrichment analysis demonstrated that CSE increased the expressions of genes related to a variety of intracellular signals, such as organic anion transport, chemical carcinogenesis, and cellular hormone metabolic processes, which were regulated by various transcription factors, including PPAR, AHR, HNF1A, ARNT, and GATA4 (Figure 1c-d). CSE-decreased genes indicated the involvement of signals, such as camera-type eye photoreceptor cell differentiation, cellular response to ammonium ion, and positive regulation of cytokine-mediated signaling pathway, although they were not significant (Figure 1e). Among them, we first focused on AHR signals, which were executed by heterodimeric transcription factors of AHR and ARNT.

### RNA-seq analysis of HepG2 cells treated with AHR agonists

To more directly observe the mechanism by which AHR signal acts on *ACE2* expression, effects of AHR agonists, such as FICZ and OMP, were evaluated in HepG2 cells. RNA-seq analysis demonstrated that FICZ and OMP increased expression of 1804 or 2106 genes, respectively, and decreased that of 830 or 1136 genes, respectively (Figure 2a). Hierarchical clustering and principal component analysis (PCA) indicated that signals regulated by FICZ and OMP were relatively more similar than they were with those regulated by CSE, but the transcriptome of FICZ differed from that of OMP, as judged from the PCA result (Figure 2b, c). The RNA-seq data suggested that the *CYP1A1* gene was strongly induced in HepG2 cells with FICZ and OMP treatments, but expression of the *ACE2* gene was clearly inhibited (Figure 2d). A Venn diagram indicated that 546 genes were commonly upregulated and 269 genes were commonly downregulated in CSE-, FICZ-, or OMP-treated cells (Figure 2e, i). Gene set enrichment analysis demonstrated that commonly regulated genes were related to metabolism of xenobiotics by cytochrome P450, which is known to be regulated by an AHR signal (Figure 2f, g, j, and k). Interestingly, an RNA-seq dataset composed of SARS-CoV-2 infection experiments (Coronasacape: a customized version of Metascape bioinformatic platform that provides a central clearinghouse for scientists to laser focus their OMICS analysis and data-mining efforts) suggested that the commonly regulated genes noted here overlapped with the genes modified after SARS-CoV-2 infection (Figure 2h, l).

**Figure 2.**
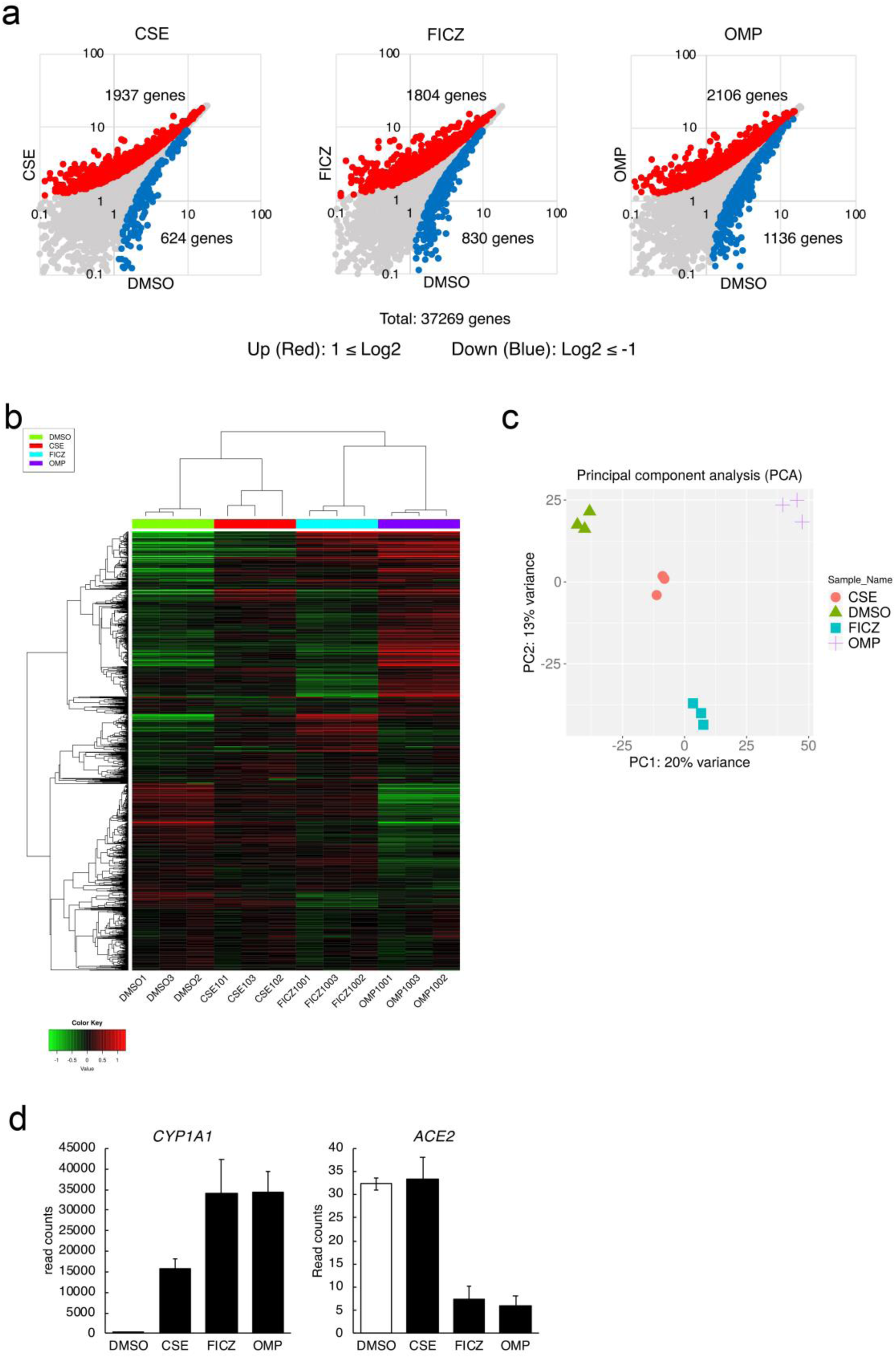
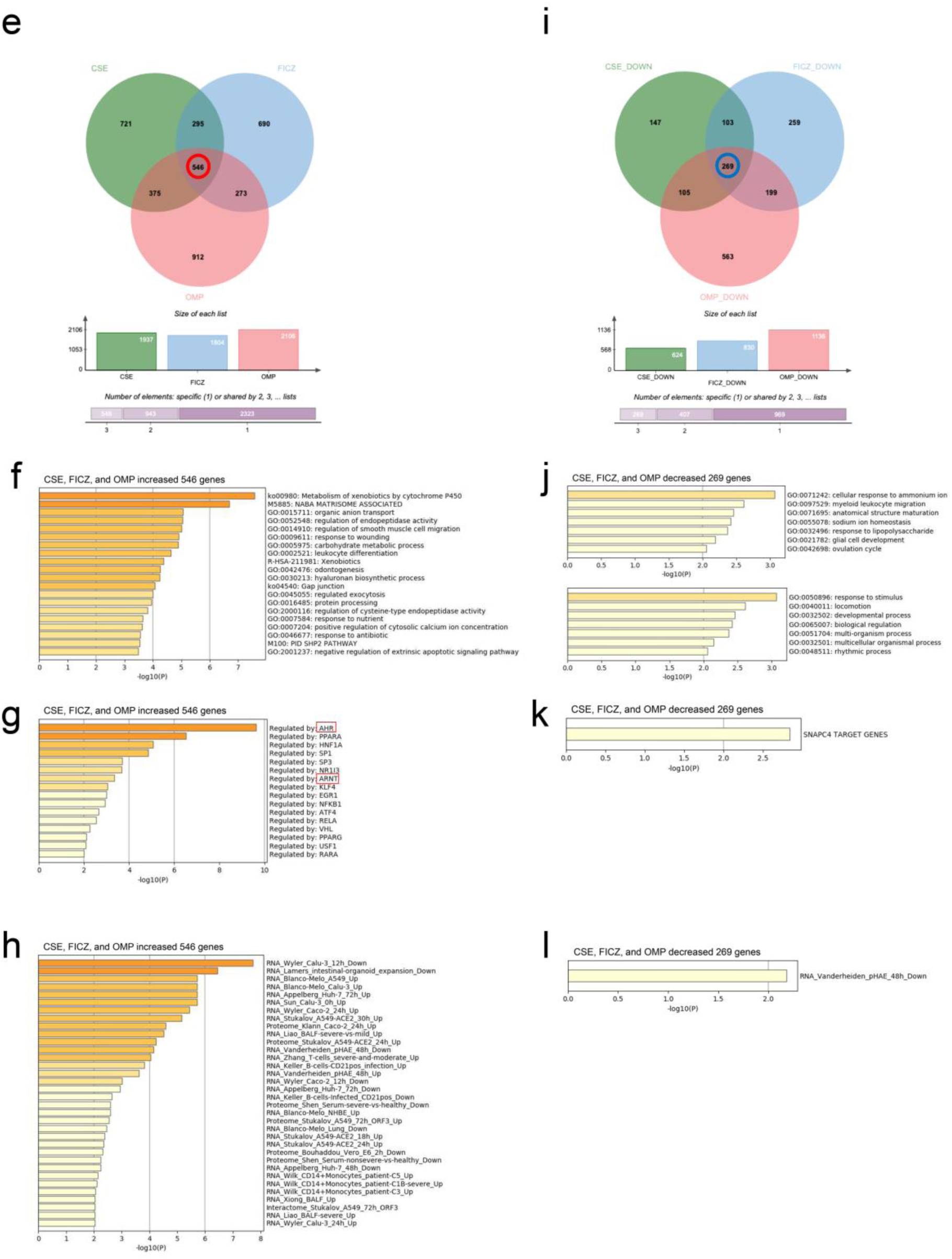
Comprehensive RNA-seq gene expression analyses of HepG2 cells treated with AHR agonists. (a) Scatter plots of gene expressions (read counts) analyzed by RNA-seq in CSE (left), FICZ (middle), and OMP (right) treated HepG2 cells are shown. Upregulated genes [log_2_(CSE, FICZ, or OMP-DMSO) ≥ 1] and downregulated genes [log_2_(CSE, FICZ, or OMP-DMSO) ≤ - 1] are indicated as red and blue dots. Hierarchical clustering (b) and principal component analysis (PCA) (c) for results of RNA-seq were performed with iDEP 0.91. Gene expression levels (read counts) of *CYP1A1* (left) and *ACE2* (right) genes in HepG2 analyzed with RNA-seq are shown. Venn diagrams of genes upregulated (e) and downregulated (i) with CSE, FICZ, or OMP treatment in HepG2 cells are shown. Upregulated (f, g) and downregulated (j, k) genes commonly regulated were enriched by GO term and TRRUST. AHR and ARNT are indicated with red boxes. Enrichment analyses of upregulated (h) and downregulated (l) genes were also performed with Coronascape (coronascape.org), a one-stop meta-analysis resource of large-scale omics data.

### Effects of AHR agonists on *ACE2* expression in HepG2 cells

qRT-PCR analysis demonstrated that FICZ and OMP efficiently induced expression of the AHR-target gene, *CYP1A1*, in a dose-dependent manner (Figure 3a). Importantly, treatment with FICZ or OMP significantly suppressed *ACE2* expression in HepG2 cells (Figure 3b). Furthermore, immunoblotting analysis confirmed that ACE2 protein expression was suppressed by treatment with FICZ or OMP, consistent with their effect on *ACE2* gene expression (Figure 3c). Analysis of the dose dependence of FICZ and OMP demonstrated that induction of *CYP1A1* and suppression of *ACE2* by FICZ both appeared to saturate at approximately 40 nM, whereas corresponding effects of OMP increased up to 100 μM (Figure 3d-g).

**Figure 3.**
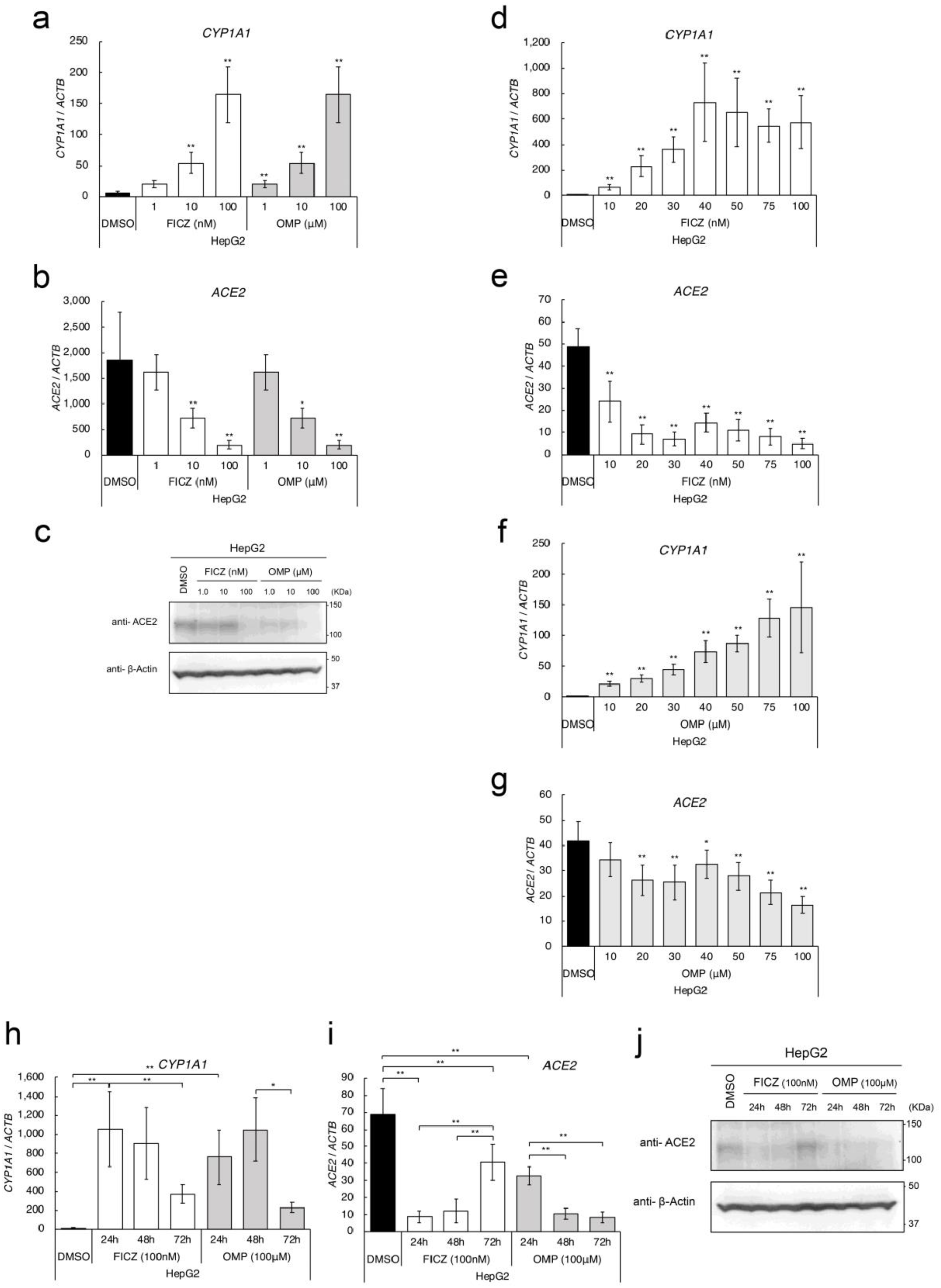
Effects of AHR agonists on *ACE2* expression in HepG2 cells. Expression levels of *CYP1A1* (a), and *ACE2* (b) genes in various concentrations of FICZ or OMP treated HepG2 cells were evaluated by quantitative RT-PCR. (c) Protein expression levels of ACE2 in various concentrations of FICZ or OMP treated HepG2 cells were evaluated by immunoblotting analysis (n=3). A representative result is shown. Dose-dependent regulation of *CYP1A1* (d, f) and *ACE2* (e, g) gene expressions with FICZ or OMP treatments were evaluated by quantitative RT-PCR. Time-dependent regulation of *CYP1A1* (h) and *ACE2* (i) gene expressions with 100 nM of FICZ or 100 μM of OMP treatments were evaluated by quantitative RT-PCR. (j) Protein expression levels of ACE2 were evaluated by immunoblotting analysis (n=3). Relative gene expression levels were calculated by using *ACTB* expression as the denominator for each cell line (n = 3). For all quantitative values, the average and SD are shown. Statistical significance was calculated for the indicated paired samples with * representing *P* < 0.05 and ** *P* < 0.01.

Next, time-dependency of AHR agonists was evaluated. Treatment with 100 nM of FICZ increased *CYP1A1* expression to a maximum at 24 hours, after which it decreased up to 72 hours, whereas *ACE2* expression was suppressed to the greatest extent at 24 hours followed by a return to the original expression level at 72 hours (Figure 3h, i). Treatment with 100 μM OMP produced delayed responses in comparison with FICZ, the highest *CYP1A1* induction and greatest *ACE2* suppression being at 48 hours (Figure 3h, i). These observations were also confirmed by protein levels with immunoblotting analysis (Figure 3j).

### Role of AHR in *ACE2* suppression in HepG2 cells

To confirm that the effects of FICZ and OMP on *ACE2* suppression were actually dependent on AHR, knock-down experiments were performed. Effective knock-down of *AHR* in DMSO or FICZ treated HepG2 cells was confirmed with qRT-PCR, and *CYP1A1* induction with FICZ treatment was significantly inhibited in AHR knocked-down cells (Figure 4a, b). Under this condition, *ACE2* expression in AHR knocked-down cells after treatment with FICZ increased up to the level in control cells, indicating a critical role of AHR in FICZ-induced ACE2 suppression (Figure 4c).

**Figure 4.**
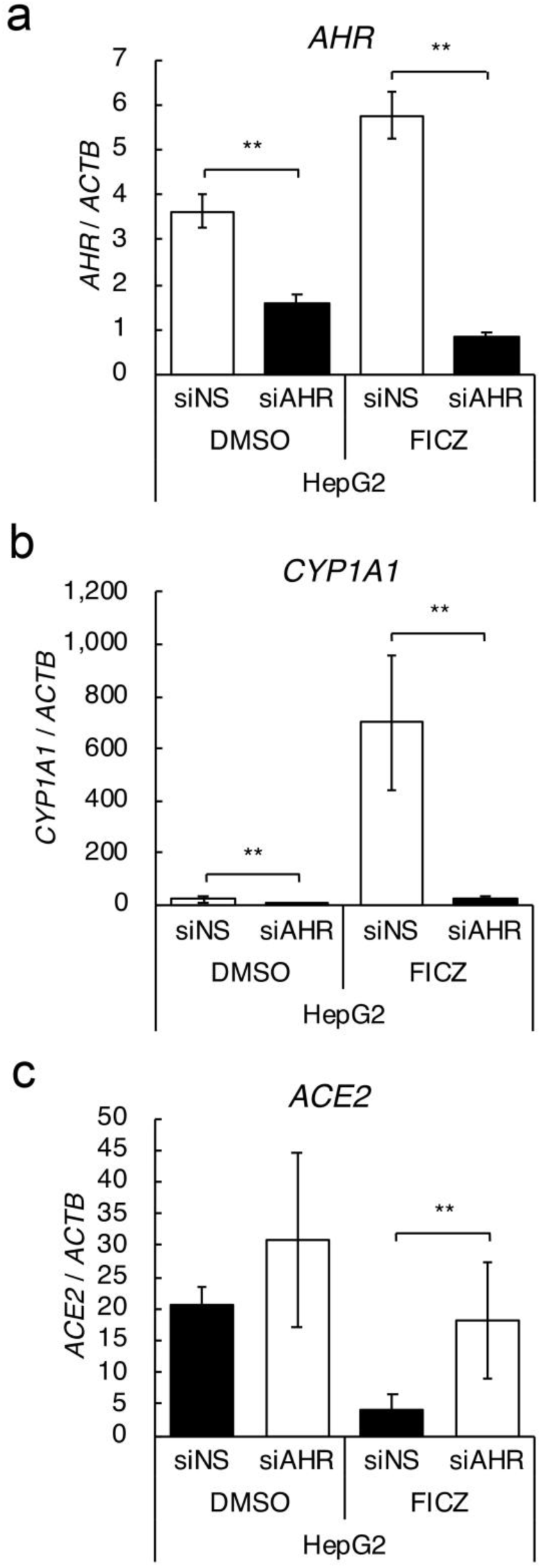
Roles of AHR on *ACE2* expression in HepG2 cells. Knock-down experiments were performed with specific siRNA for AHR. Expression levels of *AHR* (a), *CYP1A1* (b), and *ACE2* (c) genes in 100 nM of FICZ treated HepG2 cells were evaluated by quantitative RT-PCR. Relative gene expression level was calculated by using *ACTB* expression as the denominator for each cell line (n = 3). For all quantitative values, the average and SD are shown. Statistical significance was calculated for the indicated paired samples with ** representing *P* < 0.01.

### Effects of AHR agonists on SARS-CoV-2 infection of Vero E6 cells

To clarify the effect of AHR agonist-induced ACE2 suppression on SARS-CoV-2 infection of mammalian cells, Vero E6 cells expressing TMPRSS2, which are known to be susceptible to SARS-CoV-2 infection, were employed in infection experiments (Matsuyama et al., 2020). *ACE2* expression in Vero E6 cells was first evaluated by qRT-PCR and found to be downregulated by treatment with FICZ or OMP (Figure 5a). Immunoblotting analysis with antibody against a specific SARS-CoV-2 protein, nucleocapsid protein, demonstrated that pre-treatment with FICZ or OMP decreased levels of nucleocapsid protein in a dose-dependent manner, suggesting a decrease in the number of viruses infecting the cells (Figure 5b).

**Figure 5.**
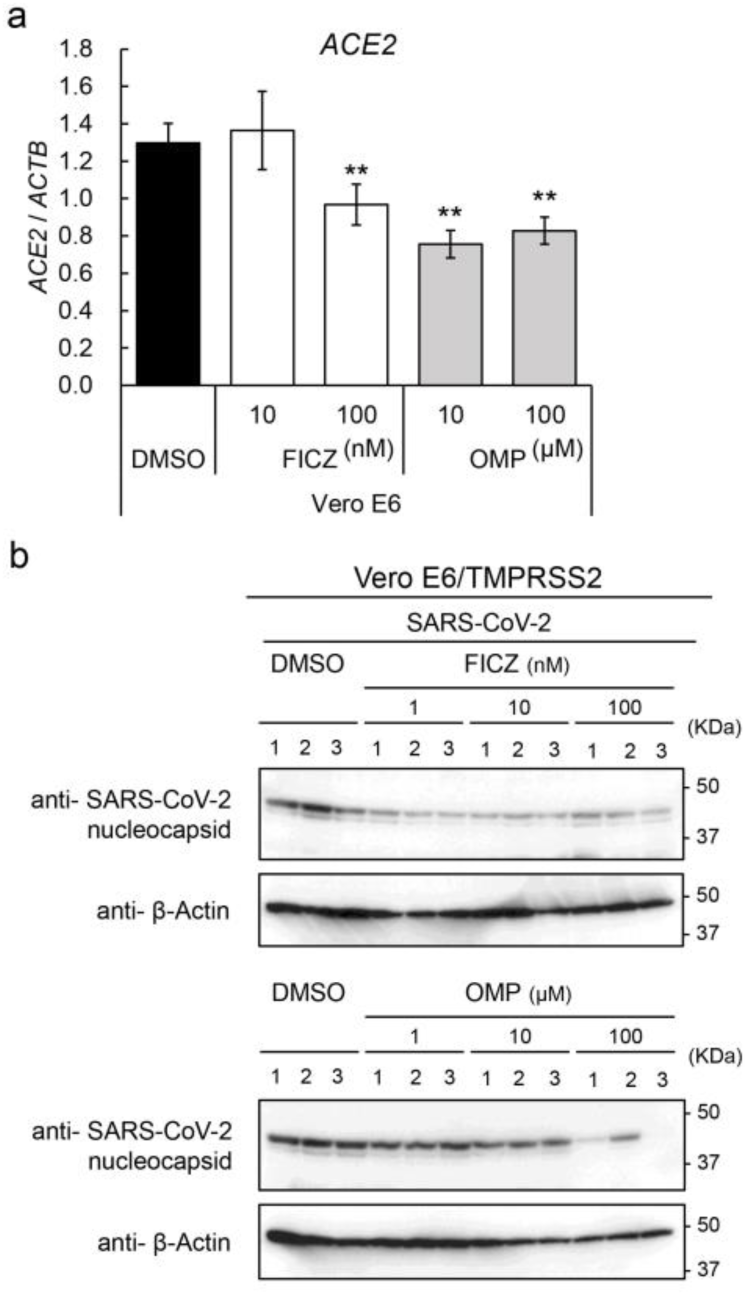
Effects of AHR agonists on SARS-CoV-2 infection to Vero E6 cells. Expression level of *ACE2* (a) genes in FICZ or OMP treated TMPRSS2 transfected Vero E6 cells was evaluated by quantitative RT-PCR. Relative gene expression level was calculated by using *ACTB* expression as the denominator for each cell line (n = 3). For all quantitative values, the average and SD are shown. Statistical significance was calculated for the indicated paired samples with ** representing *P* < 0.01. Intracellular amount of nucleocapsid protein of SARS-CoV-2 was evaluated with immunoblotting analysis (n=3).

## Discussion

ACE2 was identified as the virus receptor during the first SARS-CoV epidemic, and SARS-CoV-2 spike protein has been shown to bind ACE2 on host cells, resulting in receptor-mediated internalization of the virus (Wan et al., 2020; Zhou et al., 2020). Therefore, modulators of expression levels of ACE2 may regulate the process of SARS-CoV-2 infection. We here demonstrated that CSE, FICZ, and OMP, which could all act as agonists of AHR, suppressed the expression of ACE2 in mammalian cells, resulting in suppression of internalization of SARS-CoV-2. *ACE2* expression was significantly decreased in CSE-treated HepG2 cells, whereas *CYP1A1* expression was induced. *ACE2* expression was slightly decreased in PC9 and HSC2 cells, in which *CYP1A1* expression was slightly increased. These results suggest that *ACE2* reduction is inversely correlated with induction of one of the well-known AHR target genes, *CYP1A1*, although treatment optimization is necessary in each cell line. RNA-seq analysis of CSE treated HepG2 cells also demonstrated that CSE treatment modified a variety of intracellular signals that are regulated by the transcription factors PPAR, AHR, HNF1A, and GATA4 (Figure 1c-f). Furthermore, treatment with FICZ or OMP modified many important signals, such as response to wounding, metabolism of xenobiotics by cytochrome P450, the matrisome, and leukocyte differentiation, which were strongly suggested to be regulated by AHR (Supplementary File 2). As expected, treatment with FICZ (a tryptophan metabolite) or OMP (a proton pump inhibitor) effectively induced *CYP1A1* expression in HepG2 cells in a dose-dependent manner. In addition, both AHR agonists clearly reduced ACE2 expression at both the mRNA and protein levels. Importantly, knock-down experiments of *AHR* confirmed that reduction of *ACE2* with FICZ treatment was dependent on AHR, although details of the molecular mechanism are as yet undetermined. There are several reports of AHR-mediated suppression of gene expression. One is that AHR suppresses thymic stromal lymphopoietin (*TSLP*) gene expression via phosphorylation of p300 by Protein Kinase C (PKC) δ, resulting in decreases in acetylation and DNA binding activity of NF-κB (Jeong et al., 2019). Another is decreased expression of the AP-1 family member *Junb*, which was substantially upregulated in the inflamed skin of *Ahr*-deficient mice (Di Meglio et al., 2014). Finally, AHR was further demonstrated to form a complex with Stat1 and NF-κB in macrophages stimulated by lipopolysaccharide (LPS), resulting in inhibition of promoter activity of the *IL6* gene (Kimura et al., 2009). Since Genome Browser notes that ChIP-seq datasets in ENCODE actually indicate p300 and JUN binding on the human *ACE2* gene, ACE2 downregulation by AHR agonist treatments might occur via functional interactions between AHR and those transcription factors. Furthermore, RNA-seq analysis in the present study revealed that treatment with several AHR agonists downregulated some GATA4-regulated genes in HepG2 cells, so crosstalk between AHR and GATA4 might also be a possible mechanism of *ACE2* regulation (Supplementary File 2g).

Another interesting observation in this study was the difference in time course effects of FICZ and OMP. FICZ seemed to affect *ACE2* expression early and suppression by FICZ was quickly reduced, but the effect of OMP seemed to be more gradual and suppression by OMP remained up to 72 hours. This may be because of different activities and kinetics of their metabolism. Several other tryptophan metabolites, indole 3-carbinol (I3C), indoleacetic acid (IAA), tryptamine (TA), and L-kynurenine (KYN), and proton pump inhibitors, rabeprazole sodium (RBP), lansoprazole (LSP), and tenatoprazole (TNP), were also tested as to whether they regulate *ACE2* expression; it was demonstrated that most of them decreased expression of *ACE2* with a variety of actions and increased expression of *CYP1A1* (Supplementary File 3a, b). Similar differences were also observed among cell lines tested here (Figure 1). Optimization of each compound will be necessary in the next steps (*in vivo* experiments and clinical studies), although most of the tryptophan metabolites and proton pump inhibitors seem to be safely applicable to clinical therapeutic research. One limitation of this strategy might be that these drugs do not target the SARS-CoV-2 virus itself but just modify cellular susceptibility to it. Combination therapies of AHR agonists with anti-virus drugs, such as favipiravir or remdesivir, are therefore possible strategies for clinical application.

In addition to modifying cellular susceptibility to SARS-CoV-2, treatment with AHR agonists might stimulate immune response in the treated cells without virus infection. An RNA-seq dataset composed of SARS-CoV-2 infection experiments suggested that genes regulated by CSE, FICZ, and OMP overlapped with genes modified by SARS-CoV-2 infection, suggesting that stimulation of the immune system is involved (Figure 2h and k, Supplementary Figures 2d and h). These observations might signify a potentially useful clinical application, although further investigation will be necessary to elucidate the details.

In conclusion, we here demonstrated that treatment with CSE or AHR agonists decreased expression of ACE2 in mammalian cells, resulting in suppression of SARS-CoV-2 infection (Figure 6). Application of these compounds in clinical research and in clinical practice might be warranted.

**Figure 6.**
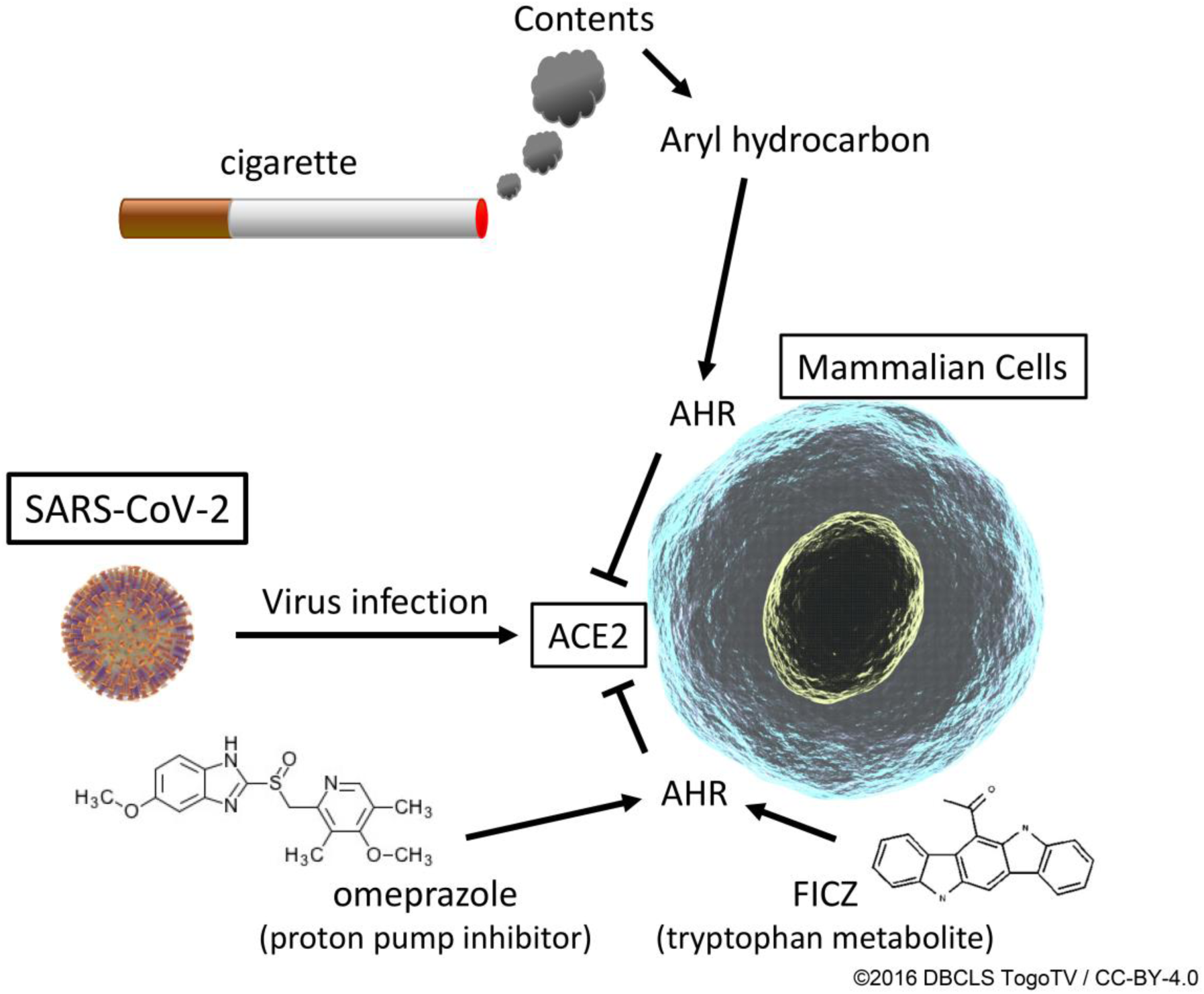
Schematic representation of effects of AHR agonists. CSE, FICZ, and OMP decreased expression of ACE2 via AHR activation, resulting in suppression of SARS-CoV-2 infection in mammalian cells.

## Materials and methods

### Chemicals

All chemicals were analytical grade and were purchased from FUJIFILM Wako Pure Chemicals, Sigma-Aldrich, or Tokyo Chemical Industry (TCI).

### CSE preparation

CSE was prepared by using a modified method reported previously (Kida et al., 2021). A detailed protocol is available at protocols.io (https://dx.doi.org/10.17504/protocols.io.bnymmfu6). Briefly, 5 filtered cigarettes were smoked consecutively through an experimental apparatus with a constant airflow (0.3 L/min) driven by a syringe, and the smoke was bubbled through 10 mL of phosphate buffered saline (PBS). The CSE obtained was then passed through a 0.22-μm filter (Millipore). The CSE was prepared immediately before each experiment.

### Cell lines and RNA preparation

Human and monkey cell lines were purchased from the Japanese Cancer Research Resource Bank (JCRRB) and have been maintained as original stocks. The cells were normally maintained in RPMI1640 or DMEM (NACALAI TESQUE) containing 10% fetal bovine serum (FBS; BioWhittaker) and 100 μg/mL kanamycin (Sigma-Aldrich), and were used within 6 months of passage from original stocks. For this study, cells (1 x 10^6^) were seeded on 10 cm diameter dishes and incubated for 24 hours, then treated with various kinds of chemicals for the indicated periods. They were then harvested for expression analysis with RNA-seq, quantitative RT-PCR, and immunoblotting. For knock-down experiments, non-specific (siNS, No. 1027310) or *AHR* (siAHR, SI02780148) siRNA (QIAGEN, Inc.) was transfected with Lipofectamine^TM^ RNAiMAX (Thermo Fisher Scientific Inc.) into HepG2 cells (1 x 10^6^/10 cm diameter dish) for 12 hours, and then the cells were incubated with DMSO or FICZ for 24 hours. Cells were then harvested and stored at −80 °C until use. Total RNA was prepared from frozen cell pellets by using NucleoSpin^®^ RNA (MACHEREY-NAGEL) according to the manufacturer’s instructions.

### Quantitative reverse transcription-polymerase chain reaction (RT-PCR)

Two µg of total RNA extracted from each cell line was reverse-transcribed using a High-Capacity cDNA Archive^TM^ Kit (Applied Biosystems). A two-hundredth aliquot of the cDNA was subjected to qRT-PCR with primers (final concentration 200 nM each) and MGB probe (final concentration 100 nM; the Universal Probe Library [UPL], Roche Diagnostics) sets as shown in Supplementary File 4 for human *ACE2*, *CYP1A1*, *AHR*, and for primate *ACE2* and *ACTB*. Pre-Developed TaqMan^TM^ Assay Reagents (Applied Biosystems) were used for human *ACTB* as an internal control. PCR reactions were carried out with a 7500 Real-Time PCR System (Applied Biosystems) under standard conditions. Relative gene expression levels were calculated by using *ACTB* expression as the denominator for each cell line.

### RNA-sequence analysis

Total RNA was processed with a TruSeq Stranded mRNA sample prep kit (Illumina, San Diego, CA, USA) (Sumi et al., 2019). Poly(A) RNA libraries were then constructed using the TruSeq Stranded mRNA library preparation kit (Illumina) and sequenced in 100-bp paired-end reads on an Illumina NovaSeq6000 platform (Sumi et al., 2019). RNA-Seq was performed in triplicate.

RNA-seq reads were quantified by using ikra (v.1.2.2) (Hiraoka et al., 2019), an RNA-seq pipeline centered on Salmon (Patro et al., 2017). The ikra pipeline automated the RNA-seq data-analysis process, including the quality control of reads [sra-tools v.2.10.7, Trim Galore v.0.6.3 (Krueger et al., 2020) using Cutadapt v.1.9.1 (Martin, 2011)], and transcript quantification (Salmon v.0.14.0, using reference transcript sets in GENCODE release 31 for humans), and tximport v.1.6.0. These tools were used with default parameters. Count tables were imported into integrated differential expression and pathway analysis (iDEP v.0.91), an integrated web application for gene ontology (GO) analysis of RNA-seq data (http://bioinformatics.sdstate.edu/idep/) (Ge et al., 2018). Quantified transcript reads were filtered at a threshold of 0.5 counts per million (CPM) in at least one sample and transformed as log2(CPM + *c*) with EdgeR (3.28.0), with a pseudocount *c* value of 4. Gene set enrichment analysis was performed in iDEP with the fold-change values returned by DESeq2 (1.26.0). False positive rates of q < 0.05 were considered enriched and investigated further with Metascape (Zhou et al., 2019). The TRRUST method (Han et al., 2015; Han et al., 2018), which analyzes human transcriptional regulatory interactions, was applied by using the web-based Metascape, a gene annotation and analysis resource algorithm (https://metascape.org/) (Zhou et al., 2019). A Venn diagram was constructed by using jvenn, a plug-in for the jQuery javascript library (http://jvenn.toulouse.inra.fr/app/index.html).

### Virus infection analysis

To observe the efficiency of infection of mammalian cells by SARS-CoV-2, SARS-CoV-2/JP/Hiroshima-46059T/2020 and TMPRSS2 transfected Vero E6 cells were employed (Matsuyama et al., 2020). Vero E6/TMPRSS2 cells in a 24 well plate were treated with FICZ or OMP for 24 hours before virus infections. The cells were then infected with SARS-CoV-2 at an input multiplicity of infection of 0.01 and incubated for 24 hours. After the cells were washed with PBS, cell lysates were obtained by directly adding SDS-sample buffer and then subjected to immunoblotting analysis.

### Immunoblot analysis

To analyze protein expression, whole cell extracts were prepared from cultured cells with the indicated treatments as previously described (Tanimoto et al., 2000). Fifty μg of extracts was blotted onto nitrocellulose filters following SDS-polyacrylamide gel electrophoresis. Anti-ACE2 (GTX101395, GeneTex), anti-SARS-CoV/SARS-CoV-2 nucleocapsid (GTX632269, GeneTex), or anti-β-actin (A5441, Sigma-Aldrich) was used as the primary antibody, diluted 1:1000, 1:1000, or 1:5000, respectively. A 1:2000 dilution of anti-mouse or anti-rabbit IgG horseradish peroxidase conjugate (#7076, #7074, Cell Signaling TECHNOLOGY) was used as a secondary antibody. Immunocomplexes were visualized by using the enhanced chemiluminescence reagent ECL Plus (Amersham Life Science).

### Statistical Analysis

Statistical tests were based on the ANOVA test, Student’s *t* test, Dunnett’s test, or the Gemes-Howell test in SPSS Statistics version 17.0 (IBM).

## Acknowledgements

We greatly appreciate the technical support provided by Ms. Chiyo Oda. We also appreciate encouragement given by Dr. Shin-ichi Hayashi (Tohoku University) and Dr. Lorenz Poellinger (Karolinska Institutet). Illustrations of the virus and the cell in Figure 5c have been reproduced from ©2016 DBCLS TogoTV / CC-BY-4.0 (https://togotv.dbcls.jp/).

## Additional information

### Author contributions

Keiji Tanimoto, Project administration, Conceptualization, Data curation, Validation, Resource, Methodology, Investigation, Visualization, Writing – original, review, and editing; Kiichi Hirota, Data curation, Validation, Resource, Methodology, Investigation, Writing –review, and editing; Takahiro Fukazawa, Software, Formal analysis, Writing –review, and editing; Yoshiyuki Matsuo, Data curation, Validation, Resource, Methodology, Investigation; Toshihito Nomura, Methodology, Investigation; Nazmul Tanuza, Methodology, Investigation; Nobuyuki Hirohashi, Supervision, Writing –review, and editing; Hidemasa Bono, Supervision, Data curation, Validation, Writing –review, and editing; Takemasa Sakaguchi, Supervision, Resource, Methodology, Investigation, Writing –review, and editing

## Additional files

### Supplementary files

**Supplementary file 1.**
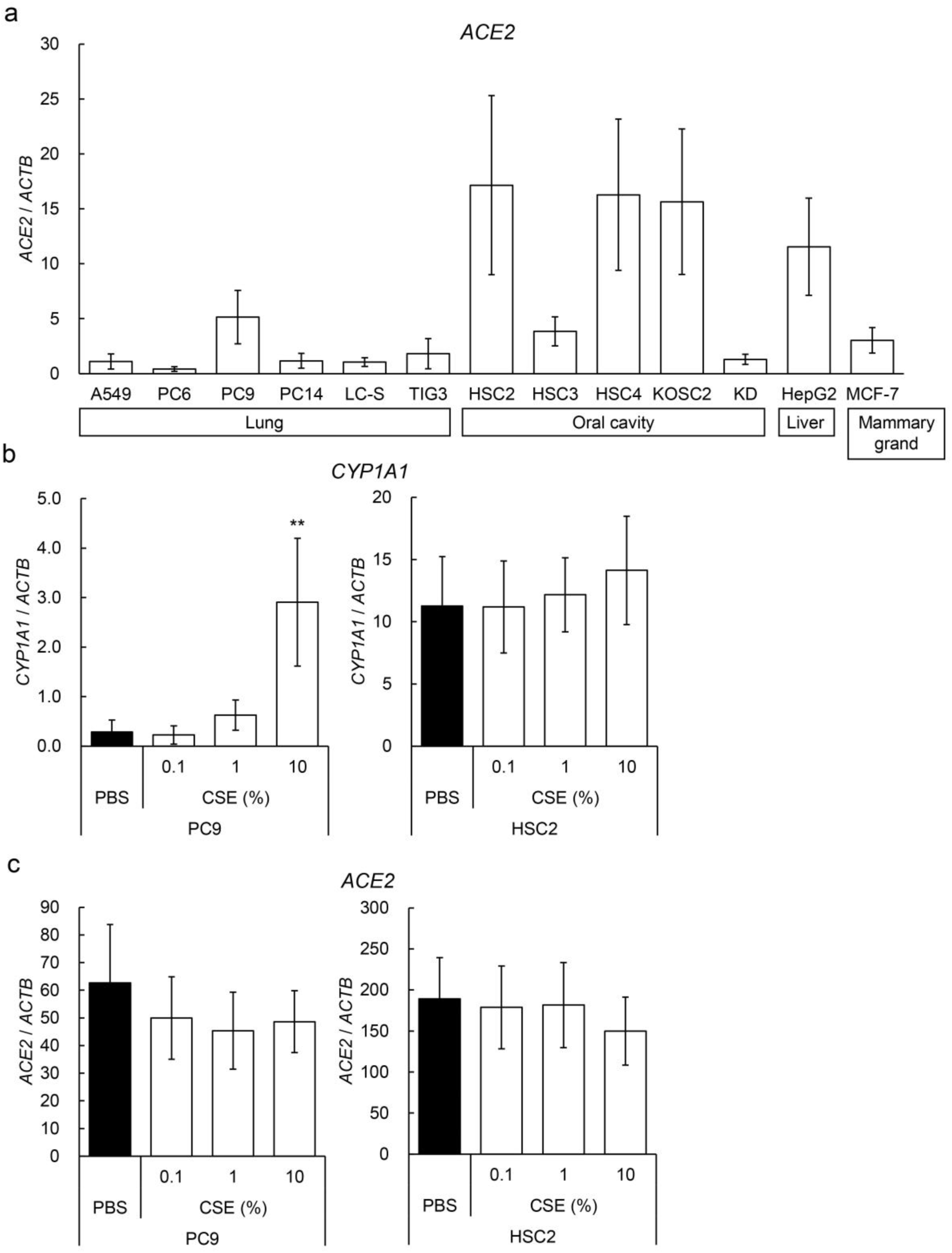
Expression levels of *ACE2* gene in various human cell lines.

**Supplementary file 2.**
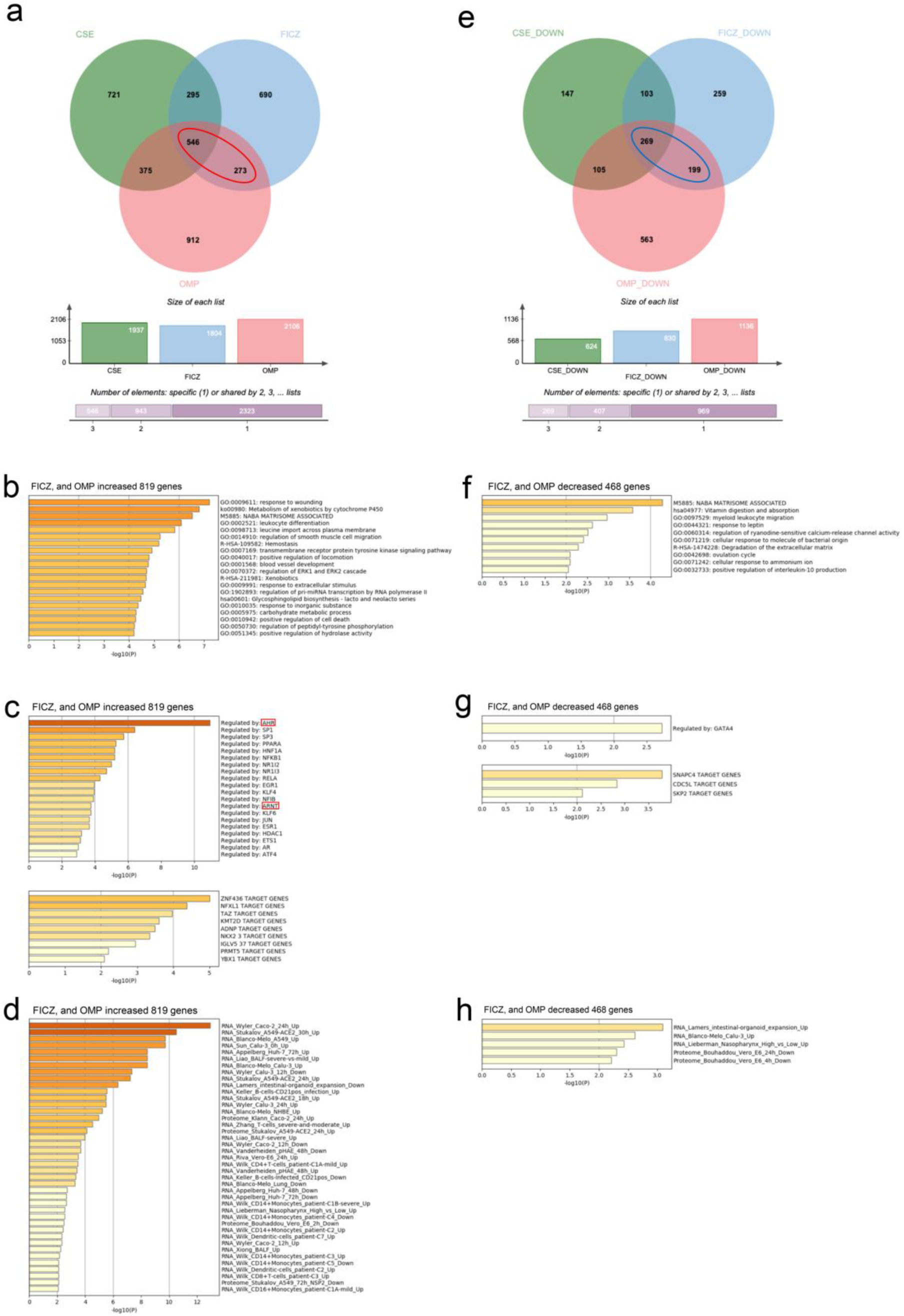
RNA-seq analyses of HepG2 cells treated with AHR agonists.

**Supplementary file 3.**
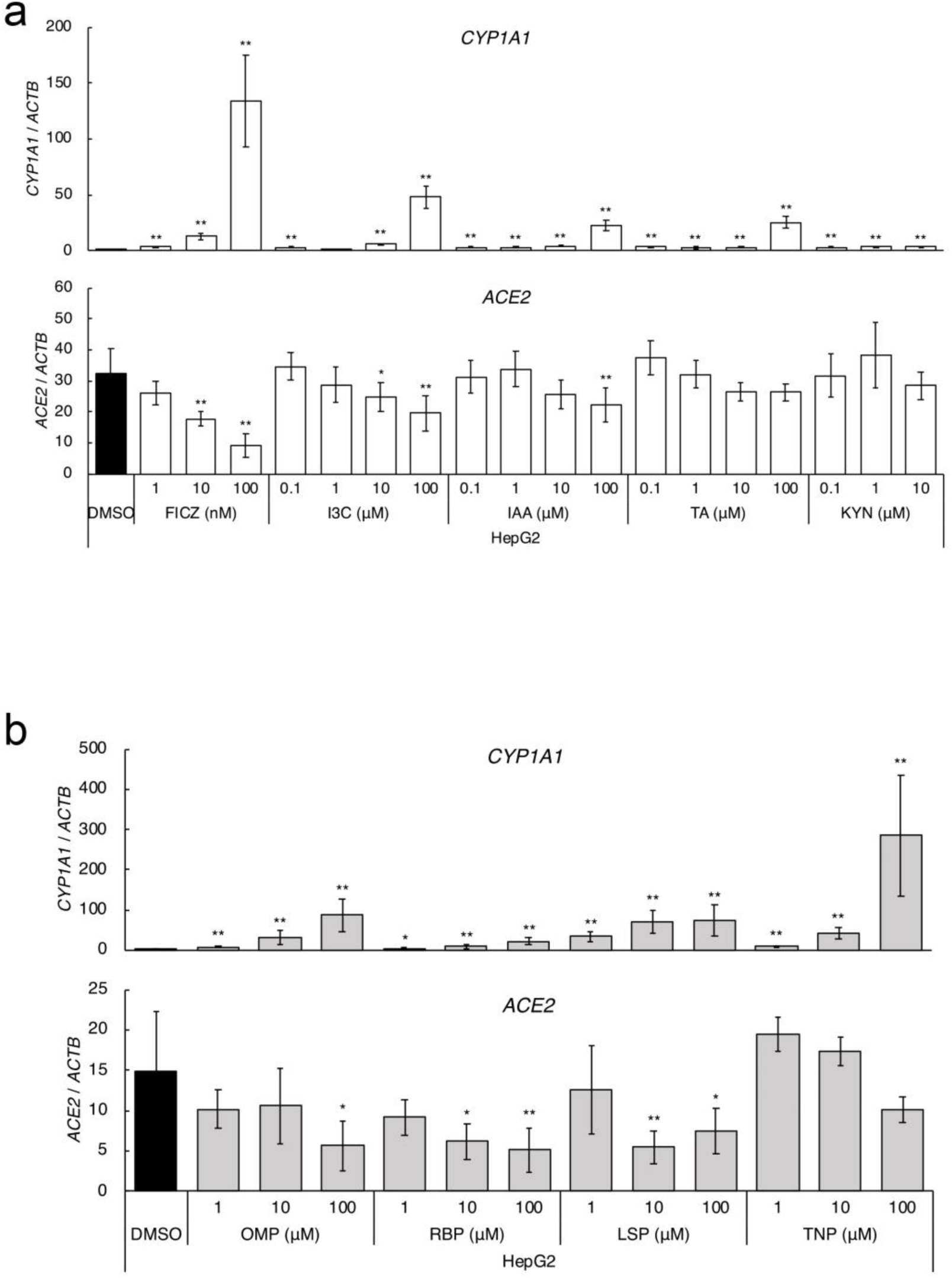
Effects of tryptophan metabolites and proton pump inhibitors on *ACE2* suppression.

**Supplementary file 4.**
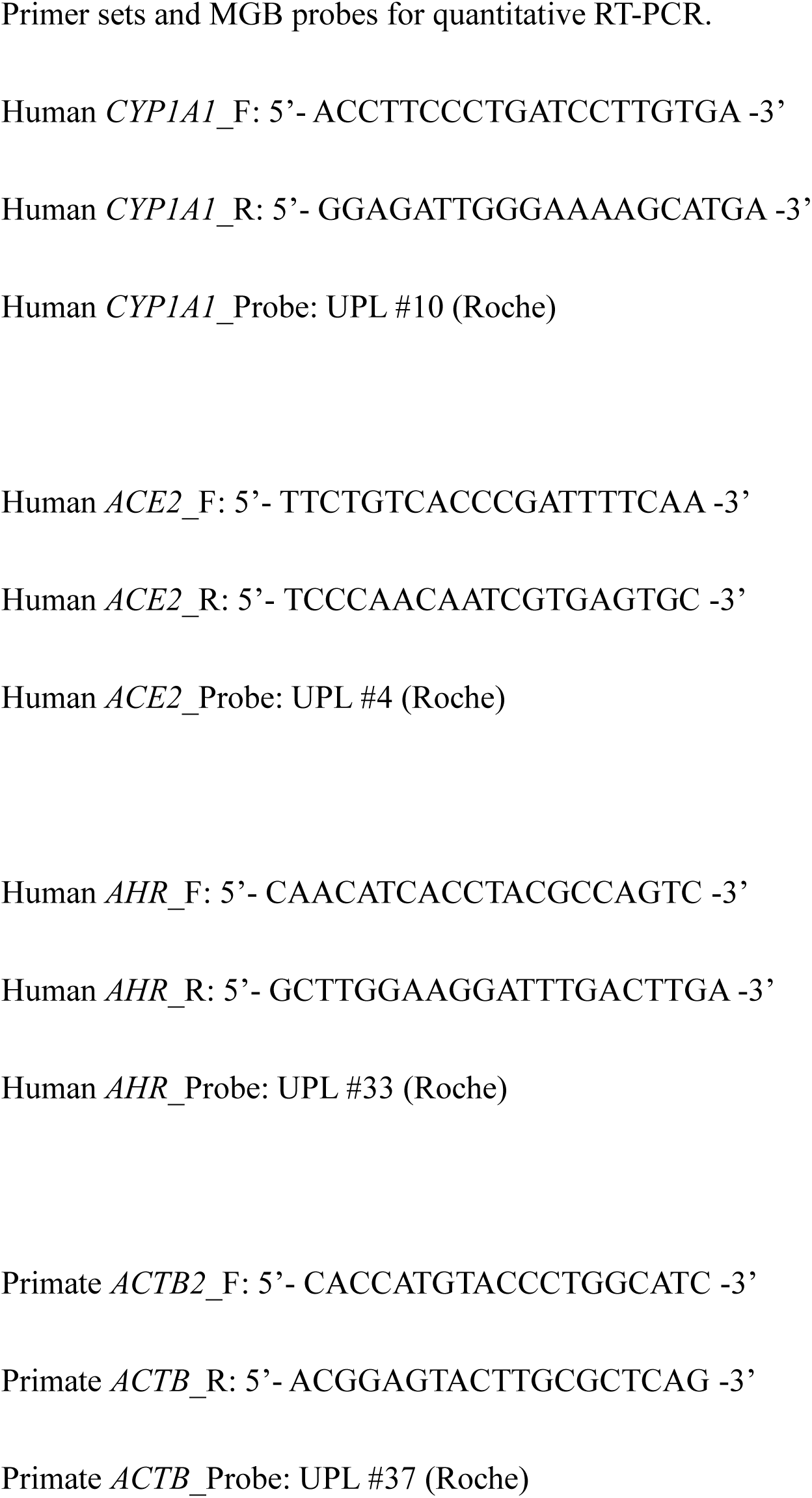
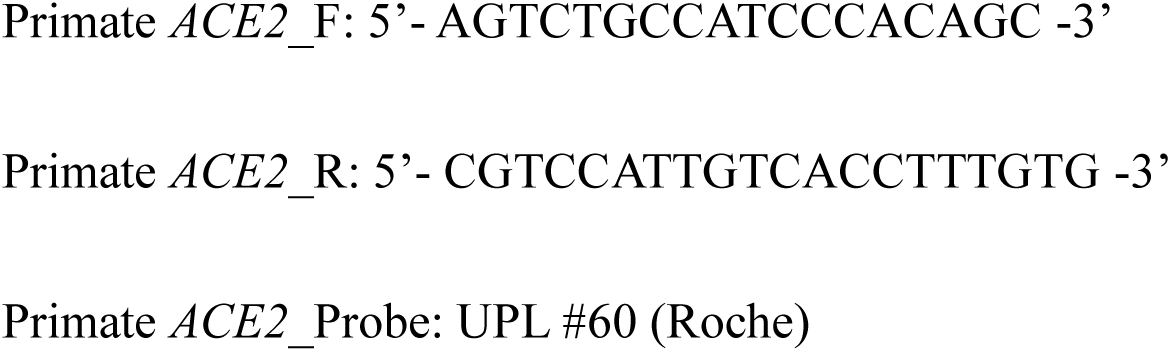
Primer and probe sequences for qRT-PCR.

## Data availability

Raw sequencing data were deposited in the DNA Data Bank of Japan Sequence Read Archive (https://www.ddbj.nig.ac.jp/dra/index-e.html; accession nos. DRR264900–DRR264911).

